# MiR-505-3p, a New Repressor of Puberty Onset in Female Mice

**DOI:** 10.1101/271718

**Authors:** Li Tong, Maochun Wang, Yuxun Zhou, Yu Li, Li Chen, Fuyi Xu, Kai Li, Junhua Xiao

**Author notes:** Corresponding author: Junhua Xiao, Yuxun Zhou Address: The College of Chemistry, Chemical Engineering & Biotechnology, Donghua University, 2999 North Renmin Road, Songjiang, 200237 Shanghai, China (cell-, To whom correspondence may be addressed. Email addresses. These authors contributed equally to this work (Li Tong, Maochun Wang).

## Abstract

Puberty onset is a complex trait regulated by multiple genetic and environmental factors. In this study, we narrowed a puberty-related QTL on chromosome X in female mice to a 1.7 Mb region and deduced that miR-505-3p was the functional gene.

In a GT1-7 cell line with stable overexpression of miR-505-3p (pGT1-7), both ribosomes and ribosome biogenesis pathways were affected, and the expression of some puberty-related genes was down-regulated. The amount of mRNA and protein products of the *Srsf1* gene decreased by 50 percent, and the expression of puberty-related genes was rescued by the overexpression of *Srsf1*. With the down regulation of *Srsf1* expression through shRNA, the mRNA accumulation of puberty-related genes decreased simultaneously in the GT1-7 cell line. The results of RIP-seq showed that SF2, the protein of the *Srsf1* gene, primarily bound ribosome protein (RP) mRNAs in GT1-7 cells.

miR-505-3p knockout female mice showed earlier vaginal opening, higher serum gonadotrophin levels and higher expression of puberty-related and *Srsf1* genes in the hypothalamus than their wild-type littermates. B6 female mice with ectopic expression of miR-505-3p in the hypothalamus showed significant growth retardance and later VO than wild types.

These results suggest that miR-505-3p may regulate puberty onset via the *Srsf1* gene and RP expression, which reveals a new regulatory pathway in mammalian puberty onset involving microRNA, SF2 and ribosome proteins.

**Author summary:** In this study, we identified miR-505-3p in a puberty-related QTL on the X chromosome as a female puberty onset repressor using a positional cloning strategy.

GT1-7 cell lines stably overexpressing miR-505-3p (pGT1-7) showed *Kiss1* and *GnRH* down-regulation. We also identified *Srsf1* as the functional target gene of miR-505-3p in GT1-7 cells. Compared to wild-type mice, miR-505-3p knockout female mice showed puberty onset four days sooner, along with the overexpression of miR-505-3p in the hypothalamus 2 days later. Thus, miR-505-3p is a new repressor of puberty onset in female mice.

## Introduction

In mammals, puberty onset is a typical complex trait defined as the transition period from the juvenile stage to adulthood during which reproductive capacity is reached ^1,2^. As a key developmental event, puberty is under the control of hypothalamic–pituitary–gonadal (HPG) axis ^3,4^, and no single molecule could be considered the sole trigger of puberty onset because puberty onset is the output of a highly coordinated operating system including thousands of genes and proteins ^5-8^. Therefore, revealing the functional genes of a complex trait is challenging ^9,10^, and genetic strategies are an effective way to reveal the mechanisms of mammalian puberty onset modulation ^2^. Mutations in *GPR54* (a G protein-coupled receptor gene) could cause autosomal recessive idiopathic hypogonadotropic hypogonadism (iHH) in humans, and the puberty-regulating function of *GPR54* has also been shown in mice with complementary genetic approaches ^10-12^. These observations suggested a crucial role of *GPR54* and its ligand *Kiss1* in the regulation of puberty. Genome-wide association studies (GWAS) identified sequence variants in or around Lin28B that were associated with age at menarche and other specific characteristics of puberty onset ^13-15^. An imprinted gene, *MKRN3,* was found to be related to central precocious puberty by whole-exome sequencing of pedigree samples ^16^. With genotype data from up to ~370,000 women, 389 independent signals for menarche age were identified by the GWAS strategy ^17^. Recently, system biology strategies have revealed that at least three gene networks that may contribute to puberty onset have been predicted ^9^. However, the ultimate mechanisms underlying puberty have not been fully elucidated.

In 2008, we began searching for puberty onset-related QTLs in mice from two inbred strains that differed significantly in puberty timing, C3H/HeJ (C3) and B6. A 9.5 Mb (ChrX: DXMit68~rs29053133) QTL on chromosome X was related to the timing of vaginal opening in female mice ^14^. We narrowed the QTL to a 1.7 Mb region by constructing 8 interval-specific congenic strains (ISCSs) and designated miR-505-3p as the candidate gene by comparing the DNA variations and gene expression levels between B6 and C3 mice.

GT1-7 cells are typically used in studies of the neuroendocrinological regulation of puberty onset due to their similarity to GnRH neurons, including the synthesis, processing, and pulsatile secretion of GnRH (gonadotropin-releasing hormone). Transcriptome analysis showed that puberty-related genes *Kiss1* and *GnRH* were inhibited in the stable miR-505-3p-overexpressing GT1-7 cell line. KEGG analysis implied that ribosomes and ribosome biogenesis in eukaryotes pathway were both affected.

*Srsf1* is a conserved regulator of pre-mRNA splicing and has been shown to be a target gene of miR-505-3p ^15^. Since its discovery, *Srsf1* has demonstrated a plethora of complex biological pathways, including several key aspects of mRNA metabolism (mRNA splicing, stability, and translation) as well as other mRNA-independent processes (miRNA processing, protein sumoylation, and the nucleolar-stress response) ^18^. Our dual luciferase assay verified that miR-505-3p could target the seed region in the 3’UTR of *Srsf1* mRNA, and western blot analysis revealed that the accumulation of SF2 protein decreased with the overexpression of miR-505-3p. Ectopic expression of *Srsf1* in the pGT1-7 cell line rescued the expression of *Kiss1* and *GnRH* genes that had been inhibited by miR-505-3p. Knocking down the expression of *Srsf1* by shRNA, we found that the expression of *Kiss1* and *GnRH* simultaneously decreased in the GT1-7 cell line. To further detect the potential regulating mechanism of *Srsf1*, we performed RNA immunoprecipitation sequencing (RIP-seq) to identify RNAs that were bound to *Srsf1*. The results revealed that the SF2 protein primarily bound to RP mRNAs in GT1-7 cells.

We also constructed a miR-505-3p knockout mouse (miR-505-3p -/-) using CRISPR/Cas9 technology and tested its VO timing and reproductive phenotypes. The miR-505-3p -/- female mice had significantly advanced VO timing (25.7 d vs 29.6 d), larger litter size, higher incidence of dystocia and more dead pups at 48 h compared to the wild-type mice. Female knockout mice also showed more lutein in the ovary; larger birth, uterus and ovary weights; and elevated serum hormone levels. qRT-PCR results showed that miR-505-3P -/- female mice had higher expression levels of the *Srsf1* in the hypothalamus compared to wild-type mice, and *Kiss1, GnRH* and *GPR54* were also highly expressed in the hypothalamus of miR-505-3P -/- female mice.

A miR-505-3p over-expression mouse model was constructed by lentivirus-mediated orthotopic injection in the hypothalamus of B6 female mice at postnatal day 15. We found significant growth retardance in LV-treated female mice compared with untreated mice, and vaginal opening in the untreated mice was observed 2 days earlier than the LV-treated mice. This result is correlated with a significant growth difference between LV-treated and untreated mice, which indicates that a lighter body weight leads to a delay in vaginal opening.

This paper presents miR-505-3p as a new regulator of puberty in female mice as determined by positional cloning, and the function of miR-505-3p is confirmed by a GT1-7 cell model, a KO mouse model and an overexpression mouse model. These results may give us insights that reveal a new regulatory pathway involving microRNA, SF2 and ribosome proteins in mammalian puberty onset.

## Results

### Fine mapping confirmed a narrow puberty-related QTL region (1.7 Mb) on chromosome X

We have identified a 9.5 Mb QTL on chromosome X that regulates puberty onset in female mice. To fine map this QTL, 8 interval-specific congenic strains (ISCSs) with intervals within this region of the B6 mouse chromosome X were substituted into a C3 background in the N7 generation. We recorded VO timing and body weights of all female mice. Nonparametric tests showed that strains 1-3, 7 and 8 had significantly different VO timing from the C3 inbred strain, not for strains 4-6 (Table. S1). Based on the QTL information and allele distribution among those strains, we ascertained a narrowed QTL between rs13483770 and rs299055848 that was approximately 1.7 Mb.

### MiR-505-3p is identified as a functional candidate gene in the QTL

There are 19 genes in the QTL region (rs13483770-rs299055848) on chromosome X, including 7 pseudogenes, 4 ncRNAs, 7 protein-coding genes, and 1 microRNA gene. To select the candidate gene, we referred to the Sanger database to search DNA sequence variations between C3 and B6 mice in this region. Data showed that sensible variations of 5 consecutive SNPs existed near the 5’ upstream region of the miR-505 gene (Table. S2). To confirm the results from the Sanger database, we sequenced the regions of interest of the 19 genes and did not find any exceptions. We compared the expression levels of these genes in the HPG axis between B6 and C3 female mice to identify differentially expressed genes. As shown in Fig. 1, except for miR-505-3p, which was expressed at a higher level in the hypothalamus of B6 mice than C3 mice, the other genes showed no significant difference between strains in either the hypothalamus or other tissues in the HPG axis (Fig. S1). Therefore, miR-505-3p was selected as the functional candidate gene in this QTL for further investigation.

**Fig. 1.**
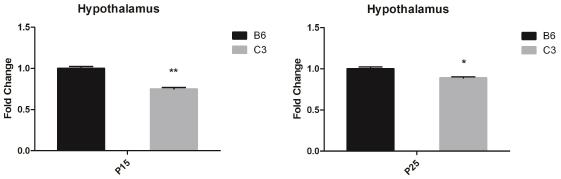
miR-505-3p mRNA levels in the hypothalamus at PND15 and PND25 in C3 mice (n = 7) normalized to B6 mice (n = 7). All values represent means ± s.e.m. (\*\**P*<0.01), and the number of mice (n) is noted here.

### Important puberty-related genes are down-regulated in a stable miR-505-3p-over-expression GT1-7 cell line

To study the function of miR-505-3p, we constructed a GT1-7 cell line with stable overexpression of miR-505-3p, and we abbreviated this cell line as pGT1-7 (transfected with pLenti6.3-miR-505-3p lentivirus). We compared the transcriptome data between pGT1-7 and the negative control group GT1-7 (transfected with pLenti6.3-nc lentivirus) and found that the expression of important puberty-related genes, *Kiss1* and *GnRH*, was lower in the pGT1-7 cell line. These results were confirmed by qRT-PCR assay (Fig. 2), implying that miR-505-3p may participate in puberty onset regulation through the inhibition of puberty-related genes.

**Fig. 2.**
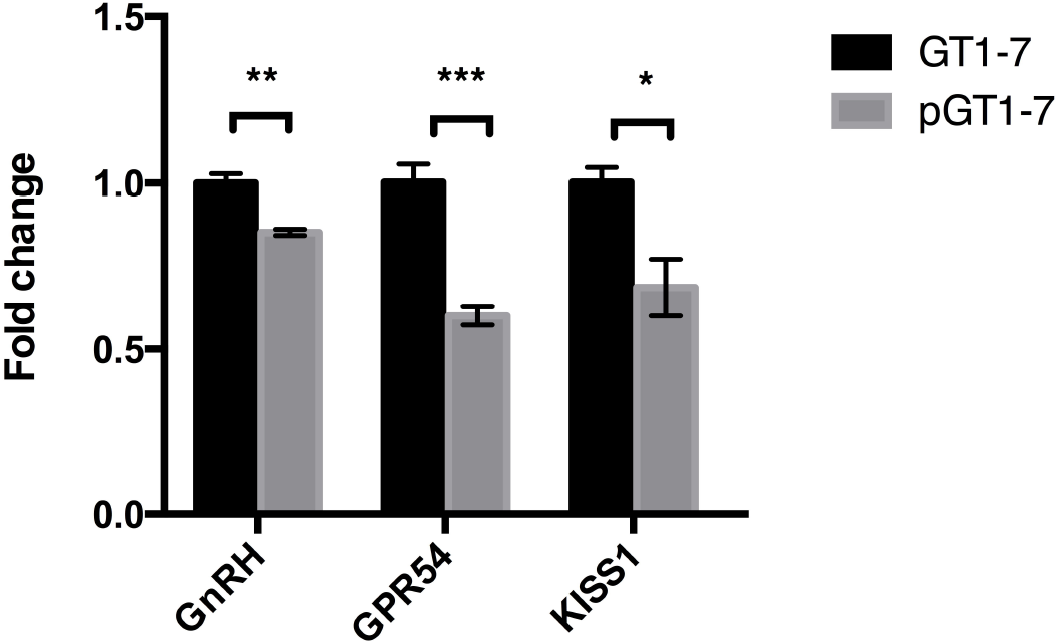
The expression level of puberty-related genes in the GT1-7 cell line that was stably transfected with miR-505-3p. Bars are means, and vertical bars represent SEM (* P<0.05, ** P<0.01, *** P<0.001).

We performed KEGG analysis on the transcriptome data to compare pGT1-7 cells with GT1-7 cells. The results show that both ribosomes and ribosome biogenesis in the eukaryotic pathway were affected in the pGT1-7 cell line (Fig. 3).

**Fig. 3.**
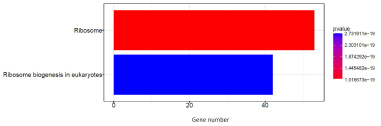
Pathway analysis of genes expressed differentially between GT1-7 and pGT1-7 cell lines.

### *Srsf1* was shown to be a target gene of miR-505-3p in the GT1-7 cell line

We verified that *Srsf1* was an effective target gene of miR-505-3p using dual luciferase reporter assay in HEK293 cells (Fig. S2). We also found that the accumulation of SF2 protein was lower in pGT1-7 cells than in GT1-7 cells.

To study the regulatory function of *Srsf1* on puberty-related genes, we over-expressed the *Srsf1* gene in pGT1-7 cells. Knocking down *Srsf1* expression in GT1-7 by shRNA simultaneously down regulated the expression of *Kiss1* and *GnRH*. When the expression of *Srsf1* was rescued, the expression level of *Kiss1* and *GnRH* increased (Fig. 4). These results indicate that miR-505-3p may inhibit the expression of puberty-related genes through *Srsf1*.

**Fig. 4.**
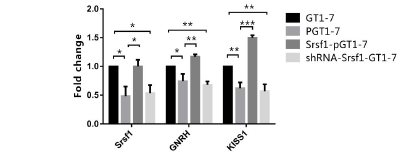
The expression levels of puberty-related genes in pGT1-7 cells after overexpression of *Srsf1*. Bars are means, and vertical bars represent SEM (* P<0.05, ** P<0.01, *** P<0.001, ns: no statistical significance)

### RIP-seq results showed that SF2 primarily bound to RP mRNAs in a GT1-7 cell line

RNA immunoprecipitation sequencing (RIP-seq) was used to identify RNAs that were bound to SF2 in a GT1-7 cell line. The SF2 protein and its bound RNAs were pulled down by an SF2-specific antibody, and the RNA component was sequenced by next-generation sequencing. We pulled down IgG protein and its bound RNAs as one control sample, and the RNA extracted from GT1-7 was another control sample. The RNA-seq data of the SF2 pull-down sample were compared to the two control samples’ data separately; the two comparison results were highly consistent, with the SF2 protein primarily binding ribosome-related mRNAs in the GT1-7 cell line (Table. 1).

**Table 1.**
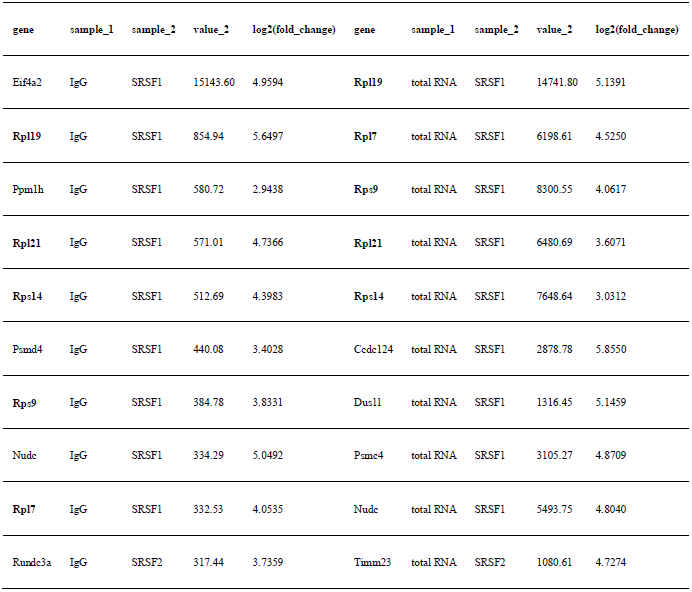
List of the top 10 genes from the SF2 pull-down sample RNA-seq data compared to two control samples’ data separately

### MiR-505-3p knockout mice showed an advanced onset of puberty and abnormal reproductive phenotypes

MiR-505-3p knockout mice were generated by CRISPR/Cas9 technology within the B6 background. In female miR-505-3p knockout mice, we observed an increased growth rate and a larger bodyweight than wild-type mice (Fig. 5A). We observed a 3.57 d and 3.87 d advance in vaginal opening in the heterozygous (miR-505-3p +/-) and homozygous (miR-505-3p -/-) knockout mice, respectively (Fig. 5B). At PND45, the mass of the reproductive system was greater in knockout mice, indicating advanced sexual development in knockout mice (Fig. 5C). *Srsf1, Kiss1* and *GnRH* expression in the hypothalamus of miR-505-3p knockout mice and wild-type mice at different postnatal days was also assessed, with the knockout mice showing higher expression levels (Fig. 6).

**Fig. 5.**
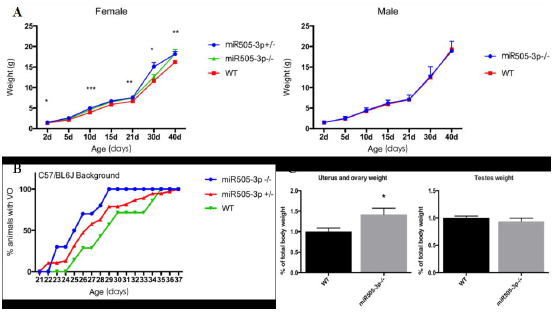
miR-505-3p KO mice show early VO.

**Fig. 6.**
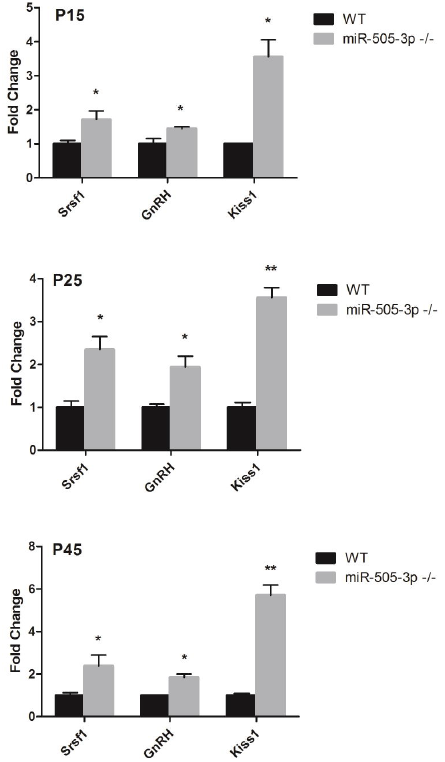
*Srsf1, Kiss1* and *GnRH* expression levels in the hypothalamus of miR-505-3p KO mice at PND15, PND25, and PND45

We then analyzed the serum levels of the pituitary gonadotropins luteinizing hormone (LH) and follicle-stimulating hormone (FSH), and the results showed a remarkable increase in female knockout mice (Fig. 7A). Moreover, the size of the litter was larger in knockout mice than wild-type mice, and heterozygous knockout mice showed more dystocia in female mice and more dead offspring at 48 h (Table. 2). Hematoxylin-eosin-stained ovarian sections of miR-505-3p knockout mice showed more corpus luteum (Fig. 7B, C). The results indicate that miR-505-3p is a repressor of puberty onset; when miR-505-3p was knocked out, puberty onset was advanced, and the reproductive system was influenced.

**Fig. 7.**
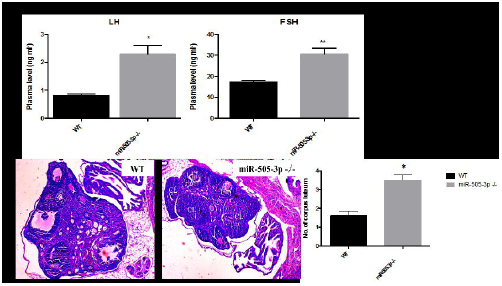
miR-505-3p KO mice phenotypes.

**Table 2.**
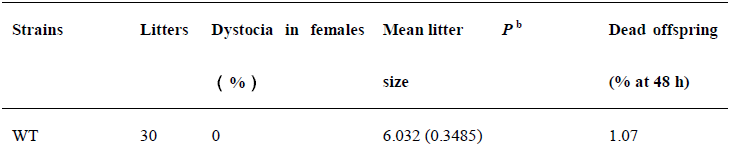
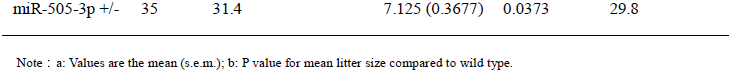
Effect of miR-505-3p deficiency on female fertility.

### Overexpression of miR-505-3p increased the time for VO compared to wild-type mice

We overexpressed miR-505-3p in the hypothalamus of B6 female mice at postnatal day 15 by lentivirus-mediated orthotopic injection. To examine the influence of miR-505-3p overexpression on puberty timing in female mice, we weighed the LV-treated female mice once every other day and found significant growth retardance in LV-treated female mice compared with the untreated mice (Fig. 8a). We observed that vaginal opening in untreated mice was 2 days earlier than in the LV-treated mice (Fig. 8b). This result correlated with a significant difference in growth between LV-treated and untreated mice, which indicates that the lighter body weight leads to a delay in vaginal opening.

**Fig. 8.**
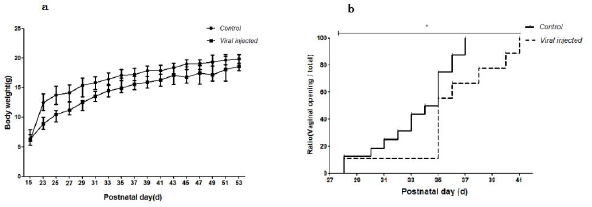
Body weight and VO timing of LV-treated female mice and control mice. a) LV-treated female mice showed significant growth retardance. b) VO timing in untreated mice was 2 days earlier than in LV-treated mice.

Moreover, LV-treated, saline-treated and untreated female mice were mated with wild-type experienced male mice to evaluate the long-term impact of hypothalamic miR-505-3p overexpression on reproduction. We found that the LV-treated female mice needed more time to procreate and had smaller newborn litters compared with control mice. The death rate of offspring before weaning was higher in LV-treated mice as well (Fig. 9).

**Fig. 9.**
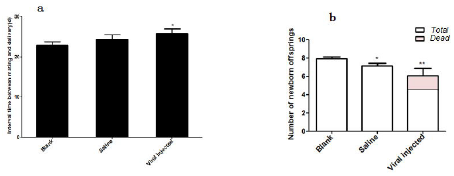
The interval between mating with male mice and death rate of offspring before weaning of LV-treated, saline-treated and untreated female mice.

## Discussion

Genetic strategies, such as positional cloning and association studies, are powerful tools to reveal the functional genes that underlie complex traits ^19-23^. Previous studies have described several important genes and complex hierarchical gene networks (including tumor-related pathways) that may contribute to the regulation of puberty onset ^24-27^. This paper aims to elucidate the functional genes and mechanisms of a puberty onset-related QTL in the mouse genome.

Testing using a GT1-7 cell line revealed that miR-505-3p could inhibit puberty-related genes and that the *Srsf1* gene was the key medium of the regulatory process. *Srsf1* is the archetype member of the SR protein family of splicing regulators and has multiple functions in the cell nucleus and cytoplasm. Moreover, *Srsf1* is also a proto-oncogene in malignant cell transformation (25). Studies have shown that the splicing-factor oncoprotein SF2/ASF, the protein of the *Srsf1* gene, activates mTORC1 ^28^. However, we did not see any activation changes with mTOR in the pGT1-7 cell line with the knockdown of the *Srsf1* gene (Fig. S3). This may be due to different cells having different gene interaction backgrounds.

Previous studies have shown that SF2 could bind to ribosomal proteins and participate in mRNA stability, transport, intracellular localization, and translation ^29^. However, there is no evidence that SF2 can bind to RP mRNAs and affect its functions. Ribosomal proteins play independent key roles in the regulation of apoptosis, multidrug resistance and carcinogenesis ^30,31^; for example, RPL22 can inhibit Lin28B, and RPS7 regulates PI3K/AKT and MAPK ^32^. These results suggest that miR-505-3p may inhibit SF2 and further affect the independent roles of the ribosome in the GT1-7 cell line, and the result of this series of events is the down regulation of puberty-related genes. There is a great deal of work to do to reveal how RP affects the molecular mechanism of puberty development.

The miR-505-3p knockout mice displayed disruptions in VO timing and several reproductive phenotypes. The overexpression of miR-505-3p female mice led to delayed VO, in addition to increased body weight, length of interval between mating with male mice and death rate of offspring. This evidence strongly suggests that miR-505-3p has a strong influence on mouse puberty and reproductive systems. There is now extensive evidence to indicate that the nuclear concentrations of SF2/ASF and hnRNP A1/A1B are fundamental in regulating the expression of specific protein isoforms within the myometrium of the human uterus. The differential expression in the upper and lower uterine segments may have a primary role in defining the formation of specific myometrial protein species associated with known contractile and relaxatory properties before and during parturition ^33^. Whether the underlying mechanism that caused dystocia was the differential expression of SF2 requires further elucidation.

The ubiquitous microRNAs construct a complex and efficient post-transcription regulatory system and play important regulatory roles in many physiological processes. According to our studies, miR-505-3p is the first microRNA that acts as a puberty regulator in female mice to be found by positional cloning; while its function was revealed by cell and animal models, the underlying mechanisms require further studies.

## Material and Methods

### Fine mapping

F1 mice were obtained from B6 and C3H lines backcrossed with C3H. Male F2 mice were genotyped, and the individuals that had at least one recombination at the specific interval were chosen and then backcrossed with a female C3H mouse to obtain the N2 generation. The female N2 continued to be backcrossed with a male C3H mouse to generate the N3 generation. N3 male mice were genotyped, and those individuals who had only one recombination at the target interval were selected and continued to be backcrossed with a female C3H mouse to generate the N4 generation. N4 mice siblings were crossed until the N7 generation. Finally, the VO timing of all female mice of the N7 progenies was recorded. All modified ISCSs were verified by genotyping genetic markers on chromosome X.

### SNP database querying between C3 and B6 mice

All SNPs between C3 and B6 mice in the QTL region (rs13483770-rs299055848) on chromosome X were queried from Mouse Genome Project in the Sanger Database (http://www.sanger.ac.uk/science/data/mouse-genomes-project).

### DNA Extraction and Sequencing

DNA extraction from the tail tip was completed using a DNA Extraction Kit (Tiangen, China), following the instructions provided by the manufacturer. All gene sequence data were from the NCBI database. In addition, the primers were designed by Primer3 (http://frodo.wi.mit.edu/primer3/) and synthesized by Shanghai Sangon Biotech, Ltd. PCR was performed in a 15 μL reaction mix containing 3 μL DNA template, 200 nM of each PCR primer,0.25 mM of each dNTP, 3 nM MgCl2, 1.5 μL 10× PCR Buffer and 1 U Hot Start DNA Polymerase (Tiangen, China) covered by mineral oil. The reaction was carried out at 95°C for 15 min, followed by 95°C for 30 sec, 55°C for 0.5 min, and 72°C for 1 min for 35 cycles. The purified PCR products were sequenced by a 3730XL sequencing instrument (ABI, USA).

### qRT-PCR

Total RNA from the hypothalamus was isolated using Trizol reagent (Invitrogen, USA) according to the manufacturer’s protocol. RNA was reverse-transcribed with M-MLV Reverse Transcriptase (ThermoFisher, USA). qPCR was carried out using SYBR green detection methods and performed according to the manufacturer’s protocol (Tiangen, China). Briefly, a 20 μL reaction contained 3 μL template, 200 nM of each primer, 10 μL 2 × SYBR Green PCR master mix, 0.4 μL ROX, and water to 20 μL. The reactions were incubated in a 96-well plate at 95°C for 2 min, followed by 40 cycles at 95°C for 15 sec and then 62°C for 32 sec with an ABI 7500 Real-Time PCR System (ABI, USA). All reactions were run in triplicate and included no template control for each gene.

### Construction of a stable miR-505-3p-over-expressing GT1-7 cell line

The dsDNA sequence of miR-505-3p was cloned into the pLenti6.3/V5-DEST lentiviral expression vector (Invitrogen, USA) and subsequently packaged into a lentivirus. GT1-7 cells were seeded at 100,000 cells per well in twenty-four-well plate and cultured in DMEM medium with 10% FBS (Gibco, USA). After 4 days of transfection, cells were selected under 6 μg/ml blasticidin for 14 days. After 10 days of expansion without blasticidin, we observed fluorescence levels under microscopy.

We used the GeneChip Mouse Transcriptome Assay 1.0 (Affymetrix, USA) to obtain the transcriptome data of pGT1-7 and GT1-7, and GT1-7 was the control. Differentially expressed genes were used to perform KEGG analysis based on the Kyoto Encyclopedia of Genes and Genomes database.

### Cell culture and RIP-seq

GT1-7 cells were cultured in DMEM with 10% fetal bovine serum (FBS). We performed RIP assays followed by high-throughput sequencing in SF2 pulldowns of GT1-7 cells, with IgG pulldowns and total RNA as the controls. Briefly, after cell lysis, 10% of the sample was removed and stored at −80°C until RNA purification was performed. The input sample was used to calculate RIP yields, and it may also be used to evaluate the quality of the RNA. RNA binding SF2 protein-specific antibody or negative control IgG was added to the supernatant. Samples were incubated with rotation at 4°C overnight. Washed protein A+G agarose was added to each IP sample, samples were incubated with rotation at 4ºC for 2 hours, and samples were washed to remove unbound material and supernatants. RNAs bound to the RNA-binding protein were purified with Trizol and sequenced by NGS.

### *Srsf1* knockdown

The expression of the *Srsf1* gene was knocked down using RNA interference (RNAi) targeting on *Srsf1* mRNA. The target sequences of shRNA for *Srsf1* (5’-GCCCAGAAGTCCAAGTTAT-3’) and non-specific shRNA (5’-TTCTCCGAACGTGTCACGT-3’) were synthesized by Obio Technology (Shanghai, China). The dsDNA sequence was cloned into the PLKD-CMV-R&PR-U6-shRNA lentiviral expression vector and subsequently packaged into a lentivirus. Finally, we received an integrated lentivirus from Obio Technology. To inhibit *Srsf1* expression, GT1-7 cells along with lentivirus in gradients of 1, 10, or 50 MOI (multiplicity of infection) were seeded at 100,000 cells per well in twenty-four-well plates and cultured in DMEM medium with 10% FBS (Gibco, USA). After 4 days of transfection, cells were selected under puromycin (0.3 µg/ml final concentration) for 14 days. After 10 days of expansion without puromycin, we observed fluorescence levels under microscopy. Total RNA was extracted from cells, and qRT-PCR was performed.

### Generation of miR-505 knockout mice

All the experiments related to mice were approved by the Institutional Animal Care and Use Committee at Donghua University. The Cas9/sgRNA co-injection method was used. Female B6 mice were superovulated using pregnant mare serum gonadotropin (PMSG) and were injected with human chorionic gonadotropin (HCG) after 48 h. The superovulated female mice were mated with B6 stud males, and fertilized embryos were collected from their oviducts. MiR-505 sgRNAs were mixed with Cas9 mRNA, and the mixture was microinjected into the cytoplasm of the female mouse embryos at the pronuclei stage. The injected zygotes were cultured at 37°C and 5% CO _2_ in air until the blastocyst stage. Subsequently, zygotes were transferred into the uterus of each pseudopregnant B6 female. DNA was extracted from the pups’ tails (founder mice), and the mice were genotyped by PCR and sequencing. Then, male and female founder mice were mated with wild-type mice. F1 mice were obtained and genotyped. Then, heterozygote F1 mice were crossed until mice positive for mutations in miR-505 were identified, indicating the successful generation of gene knockout mouse strains. The PCR products from the tail DNA samples were identified after restriction digestion with T7 endonuclease 1 (T7E1) and resolution in a 1% agarose gel. The 20 μL PCR mixture contained 10 μL master mix (Tiangen, China), 8.2 μL ddH2O, 1 μL DNA, and 0.4 μL each of the miR-505 forward and reverse primers. The PCR amplification program was as follows: 95°C for 5 minutes; 35 cycles at 95°C for 30 seconds, 55°C for 30 seconds, and 72°C for 60 seconds; and 72°C for 7 minutes. After restriction digestion with T7E1, the amplicons were separated by 1.0% agarose gel electrophoresis.

### Generation of hypothalamic miR-505-overexpressing mice

The lentiviral vector carrying miR-505-3p was constructed in our lab, and the virions were packed by Obio Technology Co., LTD (Shanghai, China). Lentivirus delivery was performed on female mice who at postnatal d 12-15 weighed 5.2-5.6 g. After 1% sodium pentobarbital anesthesia, mice were fixed on a stereotaxic apparatus (STOELTING, USA). The hypothalamic stereotaxic coordinates were determined relative to the bregma (anteroposterior [AP]=-1.53 mm, and lateral [L]=±0.24 mm) and to the skull (dorsoventral [DV]=-5.3 mm). The injection of the lentiviral preparation (2.83E+08 IU/ml, 900 nl per sample) was at a rate of 30 μL/min. Empty virion and saline as controls were also injected to female mice under the same conditions. After injection, the mice were kept at 37°C until they recovered from anesthesia. Afterwards, the mice were returned to their mothers for further investigation.

### Assessment of pubertal timing

All animal procedures were approved by the Animal Ethics Committee of Donghua University, and all experiments were conducted in strict accordance with the National Institutes of Health Guide for the Use of Laboratory Animals. All efforts were made to minimize suffering. The animals were housed under specific pathogen-free (SPF) standard conditions with a 12 h light/dark cycle and adequate water and food. Beginning on the day of weaning, mice were examined daily from 8:00 a.m. to 11:00 a.m., and the VO timing and concurrent body weight were recorded. All females were assessed for VO timing. To ensure that no environmental or methodological changes occurred during the study that might have systematically affected the results, the progenitor strains (B6 and C3H) were periodically rebred, and the timing of VO was reassessed throughout the course of experimentation to verify that the observed VO timing data were consistent.

### Equipment and statistical analysis

All the column diagrams are analyzed by Prism 5 (GraphPad Software, Inc, USA). Electrophoretic gels are shotted by ChemiDoc XRS+ (Biorad, USA) and blots are analyzed by Image Lab (Biorad, USA). Data are presented as mean±;SEM. A comparison of groups was performed using paired students t-test. Differences were considered statistically significant for P values<0.05.

## Acknowledgments

This work was supported by grants from the National Nature Science Foundation of China (Grant no. 31371257), and the Key Project of Science and Technology Commission of Shanghai Municipality (No. 14140900502).

## Supporting Information Legends

Fig. S1. Six genes showed no significant difference between the B6 and C3 strains in the hypothalamus or other tissues in the HPG axis.

Fig. S2. *Srsf1* was an effective target gene of miR-505-3p when using a dual luciferase reporter assay in HEK293 cells.

Fig. S3. No activation changes with mTOR in pGT1-7 cells were seen following knockdown of the *Srsf1* gene.

Table S1. Comparison of age at VO between eight ISCSs and the control strain C3.

Table S2. Sensible variations of 5 consecutive SNPs exist near the 5’ upstream region of the miR-505 gene between B6 and C3 mice.

## Author Contributions

JX and YZ conceived this study. LT, MW, YL and LC conducted experiments. FX and KL analysed the data. MW and YZ wrote the paper. JX critically reviewed the manuscript. All authors participated in the discussion and read and approved the fnal manuscript.

